# Golgi stabilization, not its front-rear bias, is associated with EMT-enhanced fibrillar migration

**DOI:** 10.1101/308536

**Authors:** Robert J. Natividad, Mark L. Lalli, Senthil K. Muthuswamy, Anand R. Asthagiri

**Affiliations:** Department of Bioengineering, Northeastern University, Boston, MA, USA; Department of Chemical Engineering, Northeastern University, Boston, MA, USA; Department of Biology, Northeastern University, Boston, MA, USA; Beth Israel Deaconess Medical Center, Harvard Medical School, Boston, MA, USA

## Abstract

Epithelial-to-mesenchymal transition (EMT) and maturation of collagen fibrils in the tumor microenvironment play a significant role in cancer cell invasion and metastasis. Confinement along fiber-like tracks enhances cell migration. To what extent and in what manner EMT further promotes migration in a microenvironment already conducive to migration is poorly understood. Here, we show that TGFβ-mediated EMT significantly enhances migration on fiber-like micropatterned tracks of collagen, doubling migration speed and quadrupling persistence relative to untreated mammary epithelial cells. Thus, cell-intrinsic EMT and extrinsic fibrillar tracks have non-redundant effects on motility. To better understand EMT-enhanced fibrillar migration, we investigated the regulation of Golgi positioning, which is involved in front-rear polarization and persistent cell migration. Confinement along fiber-like tracks has been reported to favor posterior Golgi positioning, whereas anterior positioning is observed during 2d wound healing. While EMT also regulates cell polarity, little is known about its effect on Golgi positioning. Here, we show that EMT induces a 2:1 rearward bias in Golgi positioning; however, positional bias explains less than 5% of single-cell variability in migration speed and persistence. Meanwhile, EMT significantly stabilizes Golgi positioning. Cells that enhance migration in response to TGFβ maintain Golgi position for 3-4 fold longer than untreated counterparts, irrespective of whether the Golgi is ahead or behind the nucleus. In fact, 35% of cells that respond to TGFβ exhibit a fully-committed Golgi phenotype with the organelle either in the anterior or posterior position for over 90% of the time. Furthermore, single-cell differences in Golgi stability capture up to 30% of variations in migration speed and persistence. These results lead us to propose that the Golgi is part of a core physical scaffold that distributes cell-generated forces necessary for migration. A stable scaffold more consistently, and therefore more productively, distributes forces over time, leading to efficient migration.

## INTRODUCTION

Epithelial-to-mesenchymal transition (EMT) and maturation of the fibrillar architecture of the tumor microenvironment are highly associated with cancer progression (1–3). While EMT and fibrillar maturation are each known to enhance cell migration, little is known about how these factors interact and cooperate to promote cell motility and invasion. A better understanding of this cooperation will yield insights into how cell-intrinsic and cell-extrinsic factors conspire to advance cancer progression.

EMT involves major changes in gene expression and cell morphology, during which an epithelial cell that is ordinarily in contact with and adhered to adjacent cells transforms into a mesenchymal phenotype, acquiring an extended uniaxial morphology and enhanced migration properties (4). In cancer, EMT is induced by a number of mechanisms, including the upregulation of transcription factors, such as Snail, and by soluble ligands, such as TGFβ (5, 6).

Meanwhile, the tumor microenvironment (TME) undergoes changes of its own. The matrix is observed to stiffen, and the density and maturity of collagen fibers increases (1, 2). Cancer cells migrate along these fibers in vivo (7, 8). We and others have studied migration along fibers using long, narrow micropatterns as an in vitro model (9–15). Cells have common features on narrow micropatterns that resemble cells migrating along fibers. This includes an elongated morphology, uniaxial migration, more effective coordination between leading edge protrusion and trailing edge contraction, as well as, more efficient retraction of the cell rear.

We recently used fiber-like micropatterns to show that EMT and fibrillar environment cooperate to regulate migration response to cell-cell interactions (11, 15). When migrating nontransformed mammary epithelial cells encounter each other on fiber-like tracks, they reverse direction and move apart. Inducing EMT with TGFβ treatment changes this migration response to cell-cell contact: instead of reversing direction, cells that have undergone EMT slide past each other. Enhanced sliding enables cells to maintain longer migration paths without changes in direction, a property that is likely to enhance dispersion efficiency, especially in tumor microenvironments where the density of cells and the frequency of cell-cell encounters are high. Notably, cells that advance further into EMT slide on progressively narrower fiber-like tracks, showing that the extent of EMT and the degree of fibrillar maturation cooperate quantitatively to modulate cell-cell interactions during migration.

Here, we investigate to what extent EMT affects migration speed and persistence of individual cells in a fiber-like microenvironment. Cells confined along fiber-like micropatterns acquire uniaxial morphology and migrate significantly better than cells on 2D (13, 16, 17). Meanwhile, EMT in 2D environments also confers uniaxial morphology and enhances motility (6). Does EMT enhance migration beyond what is already achieved by spatial confinement along fibrils? Or, are EMT and fibrillar topology redundant pathways, with little cooperative effect on motility?

To the extent that EMT and fibrillar environment cooperatively regulate individual-cell migration, a potential point of coordination involves the Golgi. Subcellular positioning of the Golgi and the associated centrosome/microtubule-organizing center (MTOC) has been linked to cell polarity and directional migration (18, 19). Anterior positioning of the Golgi was first reported in 2D wound healing assays (20). From its position at the front of the cell and ahead of the nucleus, the Golgi is thought to mediate the delivery of new membrane and adhesion proteins to the leading edge (21, 22). However, in many cell types and contexts, cell migration occurs without anterior Golgi positioning. No bias in Golgi or MTOC positioning is observed among migrating Rat2 fibroblasts *in vitro* and neurons in the developing zebrafish cerebellum (23, 24), whereas T cells invading into a collagen gel exhibit posterior positioning of the MTOC (25). Posterior Golgi positioning is also observed in cells migrating on fiber-like micropatterns (12, 13).

While some of these differences in Golgi positioning can, in part, be attributed to differences among cell systems and microenvironmental context, it raises the possibility that other attributes of the Golgi, not only its subcellular position, are involved in regulating migration. Furthermore, an important consideration is the inherent variability in behaviors at the single-cell level, even for a single cell system and microenvironmental context. Not every cell in a population moves in the same manner or with the same quantitative migration properties of speed and persistence. Moreover, individual cells may vary in how the Golgi is employed, and these variations could even occur within an individual cell over time. Single-cell analysis of Golgi and cell migration dynamics are needed to better understand the cell-to-cell variability in Golgi positioning, and how these variations are related to the variability in cell migration properties.

Finally, how EMT regulates Golgi positioning at the single-cell level is unknown. In fiber-like microenvironments, both epithelial-like African green monkey kidney cells and NIH-3T3 fibroblasts have been shown to position the Golgi behind the nucleus when migrating along fiber-like micropatterns and on fibrillar collagen in 3D cell-derived matrix (12, 13). This suggests that cells undergoing EMT would continue to employ posterior Golgi positioning. However, this conclusion is based on epithelial and fibroblast cell types with disparate backgrounds, and the effect of EMT on Golgi positioning remains to be tested directly by taking a cell system, inducing EMT, and analyzing Golgi dynamics during fibrillar migration.

To elucidate the effect of EMT on fibrillar cell migration and to better understand the role of the Golgi in fibrillar migration, we conduct here single-cell, dynamical analysis of Golgi-nuclear position during the fibrillar migration of MCF-10A mammary epithelial cells that have been induced to undergo EMT by TGFβ treatment. Our data show that TGFβ-induced EMT enhances cell motility in a fiber-like context, revealing that these cell-intrinsic and cell-extrinsic factors regulate motility in a non-redundant manner. These results have implications for how EMT and fibrillar maturation work together to promote invasiveness during cancer progression. Moreover, our analysis reveals that the stability of Golgi positioning — independent of whether the Golgi is ahead or behind the nucleus — is associated with EMT-enhanced cell motility. In fact, variations in Golgi positional stability, but not a preference for anterior or posterior Golgi position, statistically captures variations in single-cell migration speed and persistence. Thus, we identify a novel way in which the Golgi is involved in cell migration — through stabilization of its position, more so than the position itself — and propose a structural/physical role for the Golgi in cell migration.

## MATERIALS AND METHODS

### Microcontact printing

SU-8 2010 (Microchem, Westbourgh, MA) was spin-coated onto 3 inch silicon test wafers (Silicon Sense, Nashua, NH). The coating was exposed to UV light using a Quintel 4000 mask aligner (Neutronix Quintel, Morgan Hill CA) through a chrome/soda lime mask with 10 μm-wide line patterns (Front Range PhotoMask, Palmer Lake, CO). Non-cross-linked portions were etched away using SU-8 developer (Microchem, Westbourgh, MA) leaving ridges and channels. PDMS 184 was mixed with cross-linker at an 8:1 ratio (Dow Corning, Midland, MI), poured over the molds and cured for at least 2 hours at 80°C.

PDMS slabs were peeled off the silicon wafer and cut to form stamps. Prior to stamping, the stamps were placed ridge side up and plasma treated (Harrick Plasma, Ithaca, NY) for 20 seconds and quickly covered in a mixture of PBS (Invitrogen, Carlsbad, CA) and 10 μg/ml rat tail collagen I (Invitrogen, Carlsbad, CA). Stamps were left to incubate at room temperature for 30 min.

Glass-bottomed dishes (Ted Pella, Redding, CA) were flame cleaned prior to vapor-depositing a monolayer of trimethoxyglycidyl (epoxy) silane (Gelest, Morrisville, PA) overnight.

Stamps were gently rinsed in DI water, blown dry with air, and immediately placed face down on silane-treated glass dishes. The stamps and surfaces were incubated together at 37°C overnight to allow free amine groups on the collagen to crosslink to the epoxy. Following stamp removal, dishes were stored dry until use.

### Cell Culture and TGFβ treatment

MCF10A cells were infected with retrovirus encoding Golgi (GM130-RFP) and nuclear (histone 2B-GFP) markers and were selected using puromycin and hygromycin, respectively. Cells were cultured by standard methods previously described(26). Briefly, cells were passed at a 1:4 ratio every three days after reaching confluence. Cultures were maintained at 37°C and 5% CO2 in standard MCF10A growth media: DMEM F12 supplemented with 5% horse serum, 1% Penicillin/Streptomycin, Hydrocortizone, insulin, EGF (Invitrogen, Carlsbad, CA), and Cholera toxin (Sigma Alderich, St. Louis, MO).

To induce EMT, MCF10A cells were treated with 20 ng/ml TGFβ (Peprotech, Rocky Hill, NJ) added to growth medium and were cultured for 12 days prior to experiments. During the 12 days, cells were passaged at the regular frequency using the aforementioned standard methods, with the exception that cells were maintained in 20 ng/ml TGFβ-containing growth medium. This protocol matches exactly that used in our earlier work to quantify morphological and protein-expression changes induced by TGFβ treatment (11).

### Imaging and image analysis

In preparation for experiments, micropatterned surfaces were treated with 0.02% Pluronic F-127 (wt/vol) (EMD Biosciences, San Diego, CA) in PBS for 30 minutes to prevent cell adhesion to non-stamped areas. Dishes were subsequently washed once with growth media and allowed to sit for 30 minutes prior to cell seeding. Just prior to seeding, the dish was rinsed once more to supply fresh media.

Cells were seeded on micropatterned, glass-bottomed dishes and were allowed to adhere for ~25 minutes. Non-adhered cells were removed by aspiration and simultaneously flushing with fresh experimental media (MCF10A growth media +/- TGFβ) in a single step to prevent the dish from drying out. Excessive rinsing was avoided to prevent damaging the Pluronic layer. Cells were allowed to fully spread in the incubator (~30 min) before mounting the dish on the microscope for imaging.

Cell migration was tracked using an LSM700 confocal microscope equipped with a heated stage and a chamber with humidity and CO2 control (Ziess, Oberkochen, Germany). Z-stacks of the fields of interest were taken every 2.5 minutes over a 14-hour period. Preparation for track analysis was conducted using post-image processing in ZEN blue (Zeiss, Oberkochen, Germany) to make 2D projections of the Z-stacks and to stitch together overlapping fields. Cell migration trajectories consisting of nuclear and Golgi centroid positions were determined in an automated fashion, with manual input as needed, using a customized MATLAB (Mathworks, Natick, MA) code that takes advantage of the contrast provided by the nuclear and Golgi markers and provides a graphical user interface described previously(27, 28). Migration properties and Golgi states were subsequently analyzed using MATLAB.

## RESULTS

### Experimental system for automated analysis of Golgi and single-cell migration dynamics in a fibrillar context

To better understand how EMT affects cell migration in a fibrillar TME, we used narrow micropatterns of collagen to model fiber-like spatial constraints and investigated the migration behavior of non-transformed MCF-10A mammary epithelial cells and counterparts induced to undergo EMT.

Collagen was covalently micropatterned into long, narrow tracks on glass-bottomed dishes. Collagen fibers in the TME have been reported to be 2-8 μm in diameter (8). Cells migrating along fibers can spread laterally on the adhesive surface around the circumference of the fiber. Therefore, in this study, we used 10 μm-wide micropatterned tracks of collagen to approximate the circumference of a collagen fiber with a diameter of 3 μm.

Meanwhile, to induce EMT and study its effects on fibrillar migration, MCF-10A cells were treated with 20 ng/ml TGFβ for 12 days. We have shown previously that this dosage and duration of exposure reduces E-cadherin expression, upregulates N-cadherin, disrupts cell-cell adhesions, and induces an extended morphology in MCF-10A cells (11) — all features consistent with progression through EMT (6). MCF-10A cells expressing histone 2B-GFP (H2B-GFP) were used to provide high-contrast images for automated segmentation of cell location and migration trajectories. In addition, to analyze the regulation of Golgi position relative to the nucleus (GPRN), cells were transduced with an expression construct encoding GM130-RFP (Golgi marker). The migration behaviors of untreated and TGFβ-treated cells expressing these two markers were recorded by time-lapse confocal microscopy, and automated segmentation and subsequent analysis of migration trajectories were performed in MATLAB.

### TGFβ-mediated EMT increases the speed and persistence of cell migration on fiber-like collagen tracks

Migration speed and persistence were calculated from the trajectories of untreated and TGFβ-treated MCF-10A cells. Migration speed was quantified by averaging the instantaneous speeds measured at every 2.5 min interval over the entire migration trajectory of the cell. Meanwhile, the persistence of migration was determined by identifying time blocks of consecutive 2.5-min intervals during which the cell moved in the same direction without stopping or turning around. The average duration of these time blocks provided a direct measure of migration persistence.

Migration speed was 2-fold greater for cells that were treated with TGFβ than those cells left untreated (Figure 1A). Untreated cells migrated with a speed of 25.2 μm/h, whereas TGFβ-treated cells moved at a speed of 51.6 μm/h. Meanwhile, TGFβ treatment quadrupled migration persistence (Figure 1B). TGFβ-treated cells migrated with a persistence of 38.7 min compared to 10.4 min for untreated cells.

**Figure 1.**
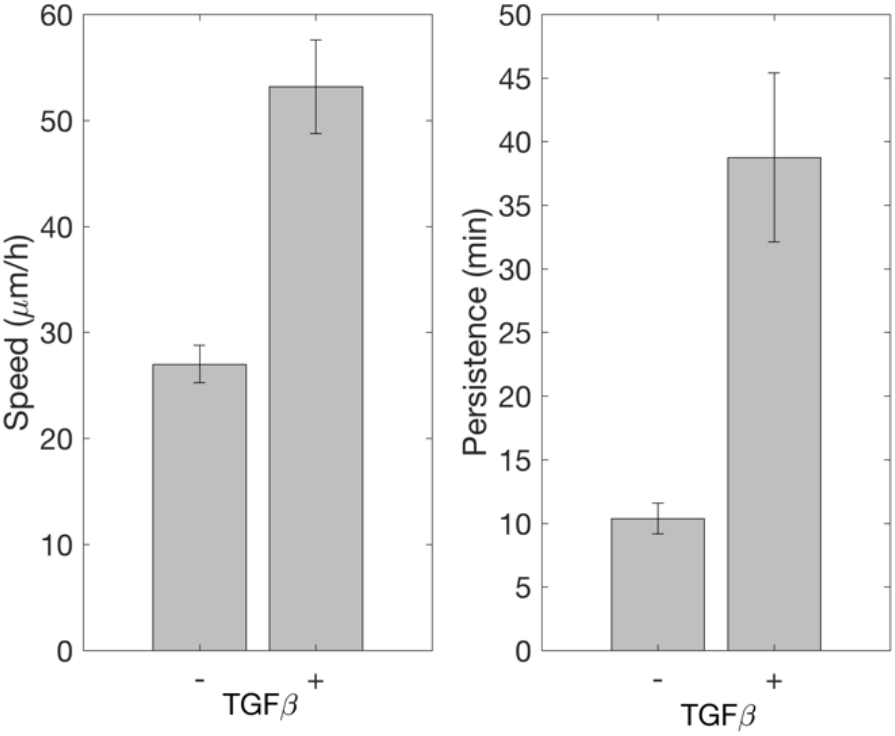
TGFβ treatment enhances speed and persistence of mammary epithelial cells migrating on fiber-like micropatterns. Migration speed (left) and persistence (right) of untreated control (-) and TGFβ-treated (+) cells are shown with error bars indicating s.e.m. with n=49 and 61 cells, respectively, across 3 and 6 independent trials, respectively. Differences in speed and persistence are statistically significant (p < 0.001, Student’s t-test).

### TGFβ-treated cells show a bias in positioning the Golgi behind the nucleus, whereas untreated cells exhibit no bias

The strong enhancement in migration, particularly persistence, suggests that TGFβ-induced EMT confers greater maintenance of front-rear cell polarity. Golgi positioning has been linked to the establishment and maintenance of cell polarity (22, 29, 30). We therefore asked whether TGFβ-induced EMT involves regulation of the Golgi position relative to the nucleus (GPRN).

To quantify GPRN, the positions of the Golgi and nucleus centroids were determined from confocal time-lapse images of H2B-GFP and GM130-RFP acquired at 2.5-min intervals (Movie S1 in the Supporting Material and Figure 2A). At each time point t, we related the orientation of the Golgi with respect to the nucleus to the direction in which the cell was migrating. The direction of cell migration was identified from the displacement of the nuclear centroid from time point *t-1* to *t*.

**Figure 2.**
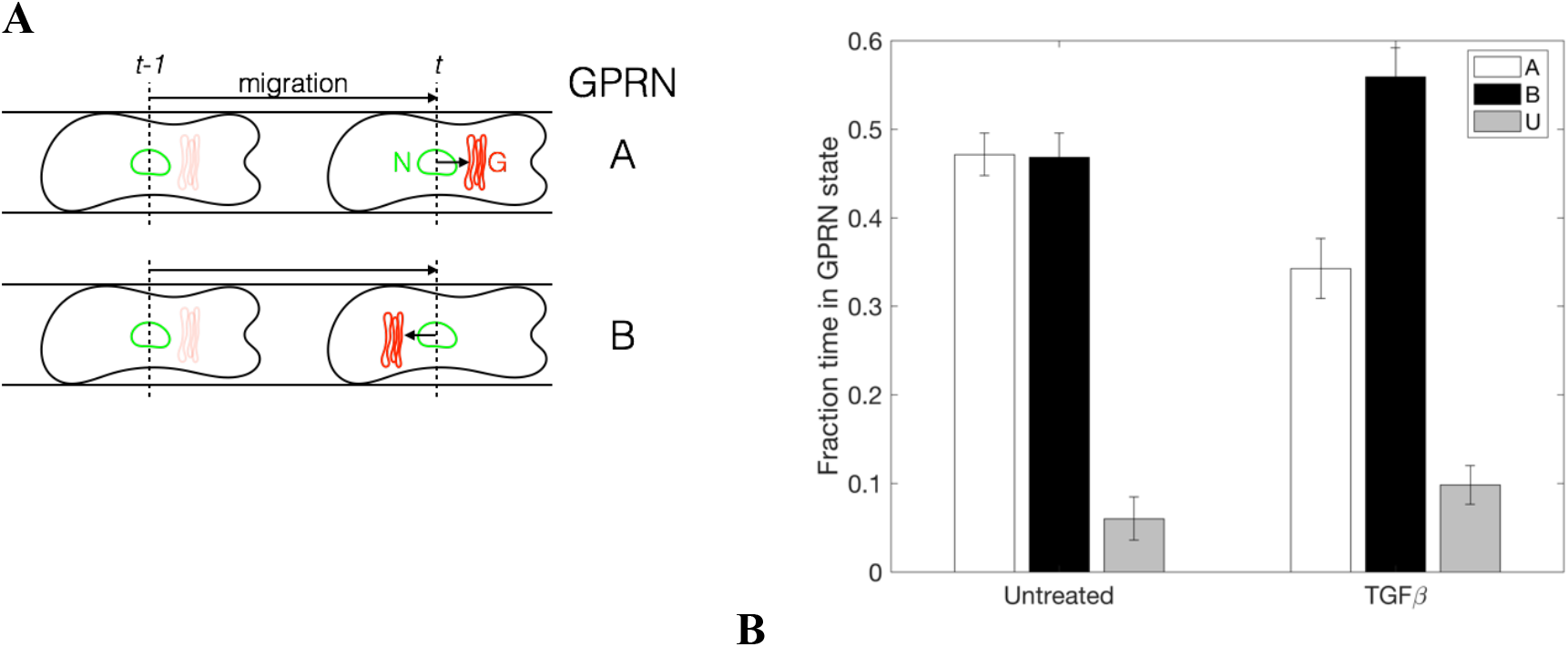
TGFβ-treated cells exhibit a bias in positioning the Golgi behind the nucleus, whereas untreated cells show an equal tendency to position the Golgi ahead or behind the nucleus. (A) Schematic of a time interval from time *t-1* to t. Direction of cell migration at time *t* is determined from the movement of the nucleus (N, green). In relation to migration direction, if Golgi (G, red) is ahead of the nucleus (top), Golgi position relative to the nucleus (GPRN) at time *t* is classified in the A state. If Golgi is behind the nucleus in relation to migration (bottom), GPRN state is B. Golgi position is unknown (U) either when the cell is not observed to move during the time interval or when the Golgi is not observable. The color of the Golgi at time *t-1* is faint to indicate that it is not considered when classifying GPRN at time t. (B) The fraction of time untreated and TGFβ-treated cells spend with the Golgi ahead (open bars) or behind (filled) the nucleus, or with the Golgi state unknown (grey), was quantified. Error bars are s.e.m., with n=49 cells (untreated) and n=61 cells (TGFβ-treated) spanning 3 and 6 independent trials, respectively.

We observed and classified three GPRN states at every time point of a cell trajectory. In the ahead state (A), the Golgi centroid was positioned in front of the nuclear centroid along the direction of cell migration. The behind state (B) occurred when the Golgi centroid was behind the nuclear centroid relative to the direction of migration. The unknown state (U) consisted of instances that could not be identified as either the A or B state. The U state occurred when the Golgi was undetectable or when the cell did not move, making the direction of migration ambiguous.

With the GPRN state recorded at every time point, a migration trajectory is represented by a string of GPRN states, such as BBBAAAUBBAABBBBUBB. This string denotes the GPRN at 2.5-min intervals as the cell migrates from its start point to its final location. In this example, state A occurs in 5 time intervals, whereas state B occurs in 11 time intervals; thus, the fractions of time spent in states A and B are 0.28 and 0.61, respectively.

To determine whether TGFβ treatment affects Golgi positioning, we quantified the fraction of time that cells spend in the three Golgi states (Figure 2B). Among untreated cells, the Golgi is observed to be ahead or behind the nucleus for an identical fraction of time of 0.47. Meanwhile, cells spend a relatively small fraction of time in the U state (0.06). These data show that untreated MCF-10A cells exhibit no preference to position the Golgi ahead or behind the nucleus during cell migration along a fibrillar track.

In contrast to untreated cells, TGFβ-treated MCF-10A cells exhibit a bias in positioning the Golgi behind the nucleus during cell migration. The fraction of time the Golgi is behind the nucleus (GPRN state B) increases to 0.56, whereas the Golgi leads the nucleus (state A) for a smaller fraction of the time (0.34). The fraction of time in the U state (0.1) is nominally higher for TGFβ-treated cells than for untreated counterparts but remains a minor state relative to GPRN states A and B. These data show that TGFβ treatment induces an apparent bias in Golgi positioning, with cells more likely to position the Golgi behind the nucleus when migrating along fibrillar tracks.

### Rearward positioning of the Golgi is specific to the subset of TGFβ-treated cells with enhanced motility

While the Golgi is, on average, more likely to be behind than in front of the nucleus among TGFβ-treated cells, it is not an absolute requirement: cells move with the Golgi ahead for a non-negligible fraction of time (0.34). Population-level averaging, however, can mask stronger relationships at the single-cell level. It is unclear to what extent individual cells differ in their migration response to TGFβ and to what extent, if any, these differences in migration response are related to the rearward bias in Golgi positioning.

To investigate the relationship between cell migration and Golgi positioning at a single-cell level, we first examined the heterogeneity of cell migration response to TGFβ treatment. Figure 3A shows the speed and persistence of individual cells. Untreated cells clustered into a relatively low-speed and low-persistence quadrant of the migration behavioral space. The cluster of untreated cells provided a reference to which the behavior of individual TGFβ-treated cells could be compared. We demarcated this reference region by determining the 90^th^ percentile values for speed (40.1 μm/h) and persistence (20.9 min) of untreated cells.

**Figure 3.**
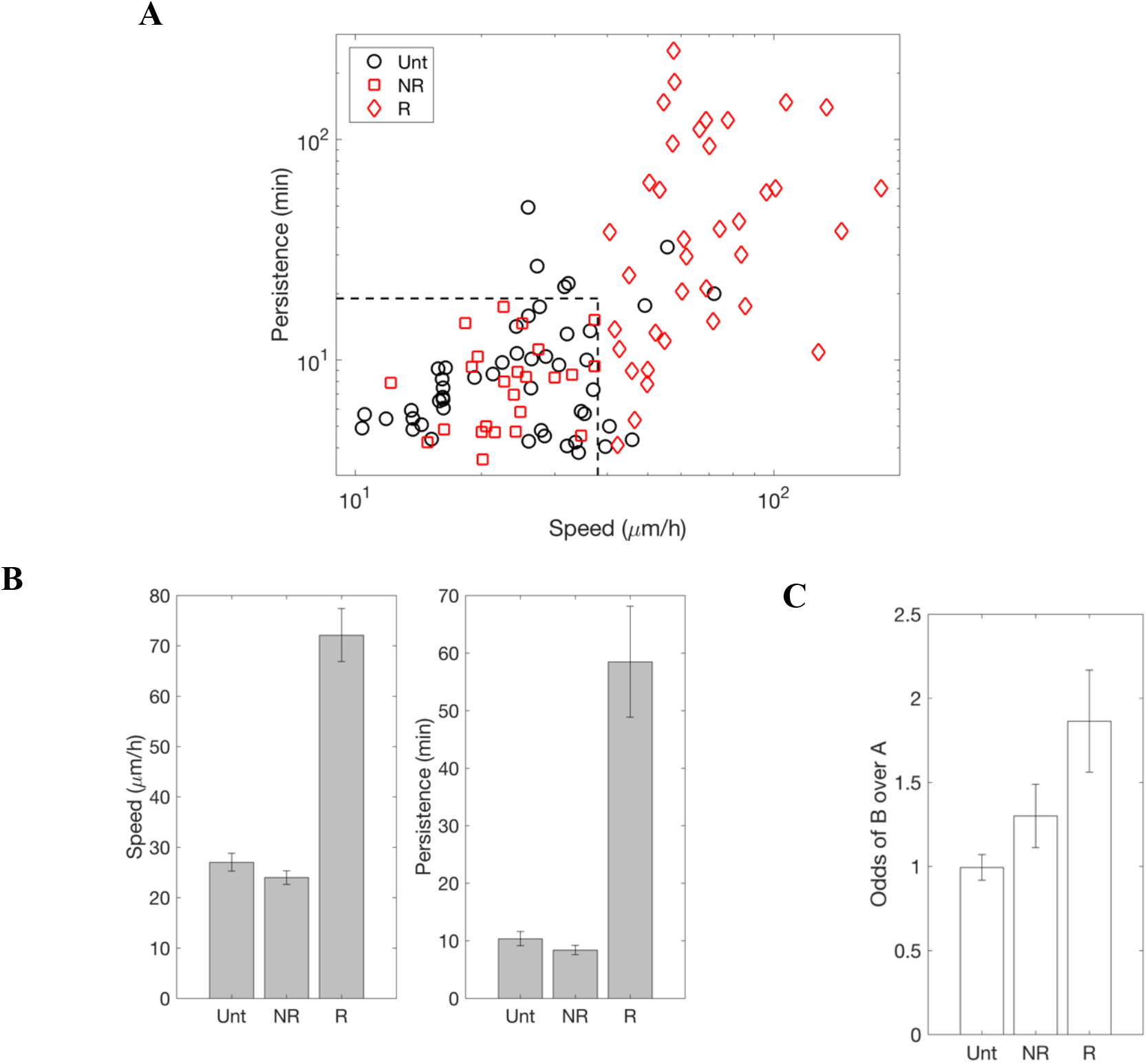
Rearward bias in Golgi position is specific to the subpopulation of TGFβ-treated cells that exhibits enhanced migration. (A) Log-log plot of migration speed and persistence for untreated cells (Unt; black, n=49) and TGFβ-treated cells (red, n=61). Dashed vertical and horizontal lines indicate 90^th^ percentile values of speed and persistence, respectively, among untreated cells. TGFβ-treated cells within demarcated region are non-responsive (NR, red squares, n=24), whereas those outside are responsive (R) (red diamonds, n=37). (B) Mean speed (left) and persistence (right) of R cells is significantly higher than that of NR and Unt cells (post-hoc Tukey’s test, p < 0.0001). Speed and persistence of NR and Unt cells are statistically indistinguishable (post-hoc Tukey’s test, p > 0.8 and p > 0.95, respectively). (C) Odds of finding a cell in state B (Golgi behind) is quantified by the ratio of fraction time spent with the Golgi behind to the fraction time spent with Golgi ahead. One-way ANOVA followed by post-hoc Tukey’s test shows that the odds of state B for R cells is statistically distinguishable from that for NR cells (p<0.05) and Unt cells (p < 0.0005), whereas odds of B for Unt and NR cells are statistically indistinguishable (p>0.8).

In contrast to untreated MCF-10A cells, TGFβ-treated cells spread out over a much broader region of the migration behavioral space. Approximately 39% of TGFβ-treated cells occupy the same low-migration quadrant as untreated cells, showing that this subpopulation did not have a motogenic response to TGFβ when compared to untreated controls. The mean speed and persistence of the non-responsive (NR) cells are 24.0 μm/h and 8.38 min (Figure 3B), respectively, comparable to 27.0 micron/h and 10.4 min (Figure 1), respectively, for untreated control cells.

In contrast to the non-responsive subgroup, the remaining 61% of TGFβ-treated cells exhibited a quantitatively stronger migration response, with increased migration speed — and in some cases, increased persistence — as compared to untreated cells. The migration speed and persistence of this responsive subpopulation is 72.1 micron/h and 58.5 min, respectively, approximately 3- and 7-fold greater, respectively, than the speed and persistence of the non-responsive cells (Figure 3B).

We next tested the hypothesis that the subpopulation of cells responsive to TGFβ exhibit a stronger bias for positioning the Golgi in the back during migration than their non-responsive counterparts and untreated cells. We quantified the odds of finding cells in GPRN state B vs A by calculating the ratio of fraction time spent in state B to that spent in state A (*f_tB_*/*f_tA_*). In untreated cells, GPRN states B and A occur to an equal extent, and the odds of finding cells in state B vs A is 1.0 (Figure 3C). The odds of finding non-responsive TGFβ-treated cells in state B is 1.3, a slight bias that is statistically indistinguishable from the no-bias behavior of untreated cells. In contrast, the odds of finding responsive TGFβ cells in GPRN state B is 1.9, a statistically-significant increase in the likelihood of finding cells with the Golgi behind the nucleus compared to untreated cells.

These results demonstrate that cells with a strong motogenic response to TGFβ are approximately two-fold more likely to position the Golgi in the back when migrating in comparison to counterparts that are untreated or do not have a motile response to TGFβ.

### A sizable fraction of TGFβ-responsive cells exhibit an “all-or-none” commitment to Golgi positioning

We noticed that even among TGFβ-responsive cells, there is a non-negligible probability (0.33), on average, of finding a cell with the Golgi ahead of the nucleus during migration on fiber-like micropatterns. This suggests that Golgi positioning in TGFβ-responsive cells is plastic, shifting ahead and behind, albeit with twice the amount of time spent with the Golgi behind than ahead of the nucleus. However, an alternate possibility is that individual TGFβ-responsive cells are not plastic and spend all their time in either state A or B, with twice as many cells fully-committed to state B than A.

Whether individual cells are plastic or are fully-committed in positioning the Golgi is unclear. To distinguish between these possibilities, we examined the distribution of fraction of time spent in states A and B among individual cells.

Among untreated cells and non-responsive, TGFβ-treated cells, the fraction of time spent in state B is symmetrically distributed and centered at 0.47 (Figure 4A). In contrast, the distribution of fraction time spent in the B state is shifted to the right among cells that have a motile response to TGFβ, consistent with a higher mean of 0.62. Interestingly, a subset of TGFβ-responsive cells (24%) spend almost all of their time (>90%) in the B state (Figure 4A, far right bin across all histograms). This full commitment to rearward Golgi positioning is not observed among the non-responsive TGFβ-treated cells or among untreated cells.

**Figure 4.**
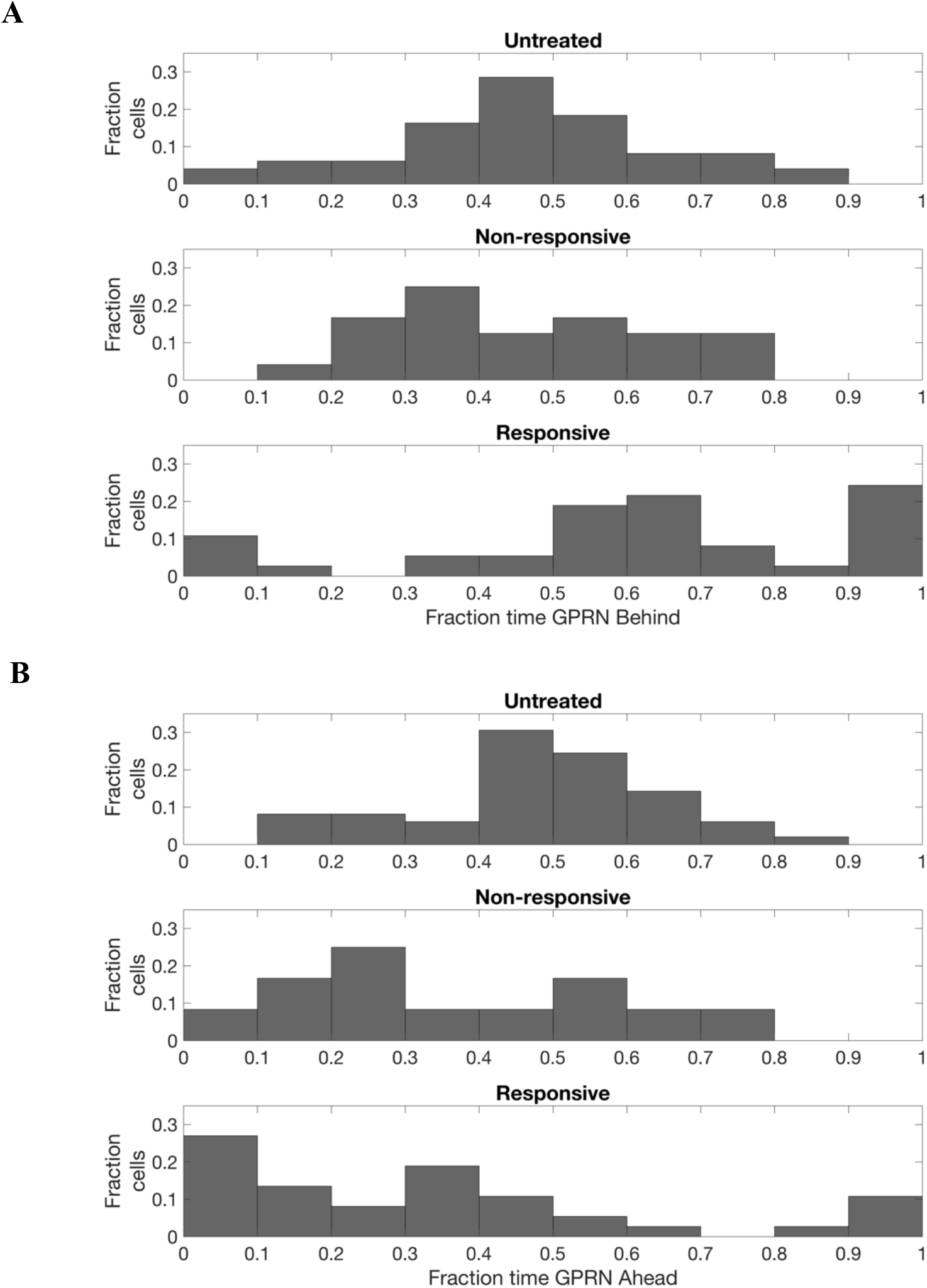
A subset of TGFβ-responsive cells exhibits all-or-none Golgi positioning and fully commits to maintain the Golgi either always ahead or always behind the nucleus. Histograms of fraction time spent with the Golgi behind (panel A) or ahead (panel B) of the nucleus are shown for untreated cells (top) and for TGFβ-treated non-responsive (middle) and responsive (bottom) cells. In both panels A and B, fully-committed cells that spend over 90% of their time with the Golgi ahead or behind the nucleus are found only in the TGFβ-responsive subpopulations.

The distribution of fraction time spent in state A (Figure 4B) shows trends similar to those observed for state B. The distribution among untreated cells is centered, with both the mean and median values of 0.47. Meanwhile, the distribution is shifted leftward for both non-responsive and responsive TGFβ-treated cells, consistent with the increased odds of finding TGFβ-treated cells in state B (Figure 3C). Notably, despite the overall reduction in the odds of finding TGFβ-treated cells in state A, a fraction of TGFβ-responsive cells (11%) spends over 90% of their time in state A (Figure 4B, far right bin across all distributions).

Taken together, this analysis reveals that a considerable 35% of TGFβ-responsive cells fully commit to maintaining a single Golgi position, either ahead or behind the nucleus, while migrating along fiber-like micropatterns. In comparison, none of the untreated or non-responsive, TGFβ-treated cells exhibit full commitment to positioning the Golgi.

These findings demonstrate that TGFβ-responsive cells have two modes of Golgi positioning during EMT-induced enhanced fibrillar migration: an all-or-none mode in which a cell commits to keeping the Golgi always ahead or behind the nucleus, and a plastic mode in which the cell shifts its Golgi between ahead and behind states. In both modes, the rearward Golgi position is the preferred state. In the all-or-none mode, a cell is twice as likely to commit to state B than A, as 24% of cells fully commit to the B state versus 11% of cells commit to the A state. In the plastic mode, cells spend more time in B than in state A.

### Golgi positioning is stabilized among TGFβ-treated cells with enhanced motility

Thirty-five percent of TGFβ-responsive cells fully commit to positioning the Golgi either at the front or the rear of the nucleus. It suggests the hypothesis that TGFβ not only increases the prevalence of state B, but also stabilizes Golgi positioning.

To test this hypothesis, we quantified the stability of Golgi positioning in individual cells. We determined the duration over which a GPRN state is contiguously maintained before it switches to another GPRN state along the migration trajectory of individual cells. A GPRN state that persists for long durations before switching to another state has a larger lifetime and greater stability than a state that is occupied for short durations.

For example, for a cell whose GPRN states at 2.5 min intervals are represented by the string BBBAAAUBBAABBBBUBB, we condense the string to B_3_A_3_U_1_B_2_A_2_B_4_U_1_B_2_, with each subscript denoting the number of time steps during which the GPRN state is maintained. For each GPRN state, averaging the associated subscript values over the entire migration trajectory provides a direct measurement of its lifetime. For the cell in this example, the average lifetime of GPRN states A, B and U are 1.5, 2.75 and 1 time steps, respectively, or 3.8, 6.9 and 2.5 min, respectively.

We determined the stability of Golgi positioning among untreated and TGFβ-treated cells. The average lifetime of states A and B are 11.4 and 18.3 min, respectively, among all TGFβ-treated cells compared to 8 min for both states among untreated cells (Figure 5). Within the TGFβ-treated population, the responsive subgroup has a stability of 15.7 and 26.6 min for the A and B states, respectively. In comparison, among the non-responsive sub-population, the stability of states A and B are 4.98 and 6.18 min, respectively (Figure 5). These results demonstrate that the stability of Golgi positioning in non-responsive, TGFβ-treated cells is on par with untreated cells. Meanwhile, cells that enhance motility in response to TGFβ exhibit a substantive stabilization of Golgi positioning, with state A being 3.1-fold and state B being 4.3-fold more stable in TGFβ-responsive cells than in non-responsive cells. Altogether, these results confirm our hypothesis that cells with a motile response to TGFβ treatment increase the stability of Golgi positioning. Furthermore, the data show that both GPRN states, A and B, are stabilized by TGFβ-treatment.

**Figure 5.**
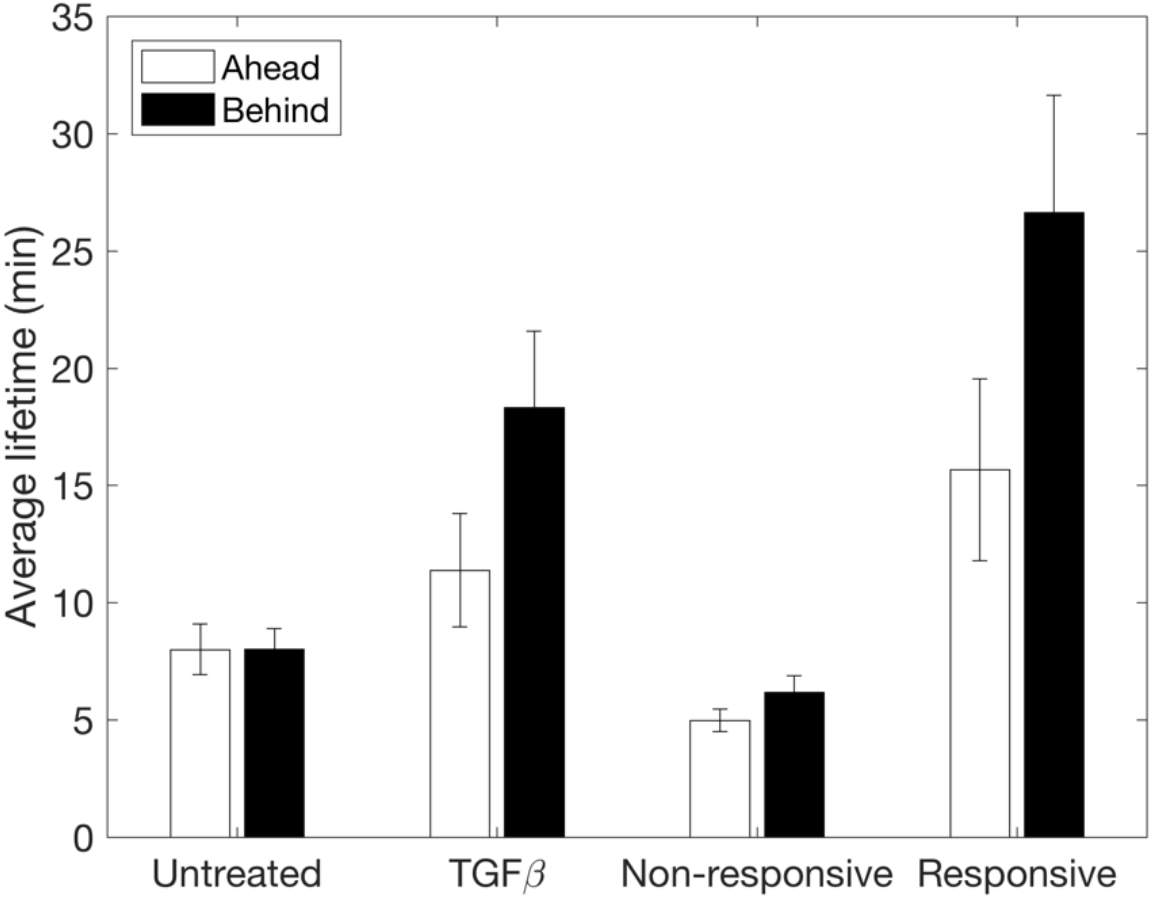
The stability of Golgi position is significantly increased among cells that enhance motility in response to TGFβ treatment. Stability of Golgi positioning was quantified as the average amount of time that cells continuously maintained the Golgi ahead (open) or behind (filled) the nucleus. The average lifetime of each Golgi position was calculated for untreated (n=49 cells) and TGFβ-treated cells (n=61 cells), and for the subpopulation of cells that were non-responsive (n=24 cells) and responsive (n=37 cells) to TGFβ treatment. Higher lifetime corresponds to greater stability.

### Golgi positioning is stabilized even among individual TGFβ-responsive cells that modulate Golgi positioning

Stabilization of Golgi positioning at the population level may occur in at least two ways at the single-cell level. One possibility is that stable Golgi positioning is unique to the 35% of TGFβ-responsive cells that exhibit full commitment to GPRN states A or B; meanwhile, the remaining plastic TGFβ-responsive cells that modulate Golgi positioning have short lifetimes in each Golgi state. Alternatively, Golgi stabilization may also occur in cells with plastic Golgi positioning: that is, although these cells divide their time between GPRN states A and B, the lifetime of one or both states may be extended.

To distinguish between these possibilities, we examined the distribution of Golgi stability among individual cells within the population. The cumulative distribution of B state stability was highly similar between untreated and non-responsive, TGFβ-treated cells (Figure 6A). In contrast, among the responsive TGFβ-treated cells, the cumulative distribution was clearly shifted to the right. In fact, the distribution for TGFβ-responsive cells deviates from non-responders after accounting for just 5% of the population, indicating that the vast majority of TGFβ-responsive cells, including those with plastic Golgi positioning, exhibit more stable GPRN states than non-responsive cells. Consistent with this high penetrance of Golgi positional stability among TGFβ-responsive cells, both the mean (26.6 min) and median (11.4 min) values of GPRN stability among the responsive cells were significantly higher than the values (6.2 and 3.2 min, respectively) for non-responders. These results show that the stability of the B state is stabilized broadly among TGFβ-responsive cells, not just the 24% of the responsive population that spends nearly all their time in the B state.

**Figure 6.**
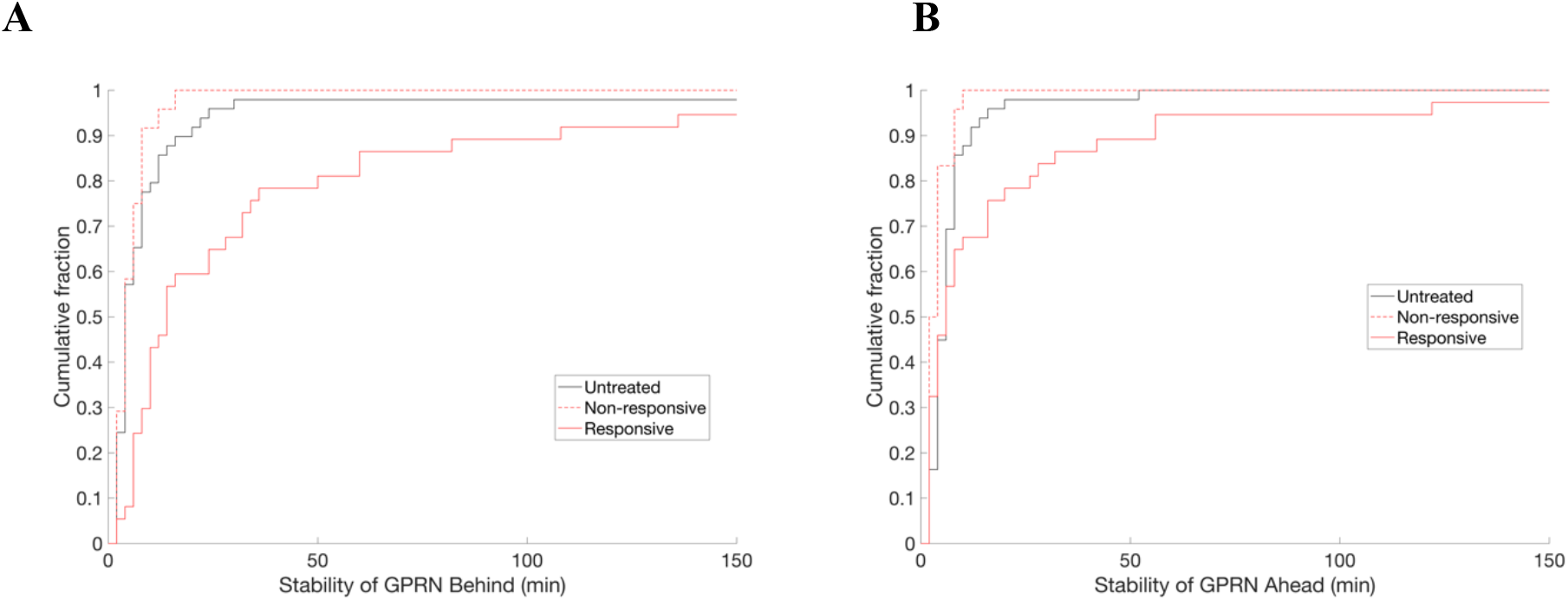
Golgi position is stabilized among a significant fraction of TGFβ-responsive cells. Cumulative histogram of the stability of the Golgi behind (panel A) or ahead (panel B) of the nucleus for untreated (black), non-responsive (red dashed) and responsive (red solid) subpopulations of TGFβ-treated cells.

The stabilization of the A state showed similar trends to the B state, albeit with quantitatively weaker effects. The cumulative distribution of the stability of the A state is similar among untreated and TGFβ-treated non-responsive cells (Figure 6B). Among TGFβ-treated responsive cells, the first-half of the distribution (below a cumulative probability of 0.5) is similar to that of untreated and non-responsive cells. Above a cumulative probability of 0.5, the distribution for responsive cells stretches rightward, extending over a higher range of stability values than for untreated and non-responsive cells. Consistent with these observations, the median value of the stability of the A state is similar for non-responsive and responsive cells at 3.2 min and 3.0 min, respectively. Thus, the higher mean stability of the A state among responsive cells (Figure 5) is attributable to approximately 50% of responsive cells for which the A state stability extends over longer duration than for non-responsive and untreated cells (Figure 6B). Thus, as with state B, state A is stabilized among a broad group of responsive cells, not merely among the ~11% of cells that spend almost all their time in the A state.

Taken together, this analysis shows that both the A and B states are stabilized not only among the responsive cells that are fully-committed to a particular Golgi state, but also among responsive cells that maintain plasticity in shifting between A and B states.

To further test the conclusion that plastic TGFβ-responsive cells that modulate between A and B states maintain those states with greater stability than untreated or non-responsive counterparts, we selected plastic cells from the TGFβ-responsive population and quantified their Golgi stability. We identified plastic cells as those cells that maintain the Golgi in a single position at most for a specified maximum fraction of time. By varying the maximum fraction of time between 0.55 and 0.95, we tuned the stringency of the plasticity criterion. For low values (e.g., 0.55), cells at most could spend 55% of time in a single Golgi state in order to be counted as plastic. Under these stringent conditions, few TGFβ-responsive cells (n=5) met the criterion (Figure 7). In contrast, when the plasticity criterion is relaxed to allow cells that spend as much as 95% of their time in a single Golgi state to still count as being plastic, more TGFβ-responsive cells (n=30) qualify as having plastic Golgi positioning.

**Figure 7.**
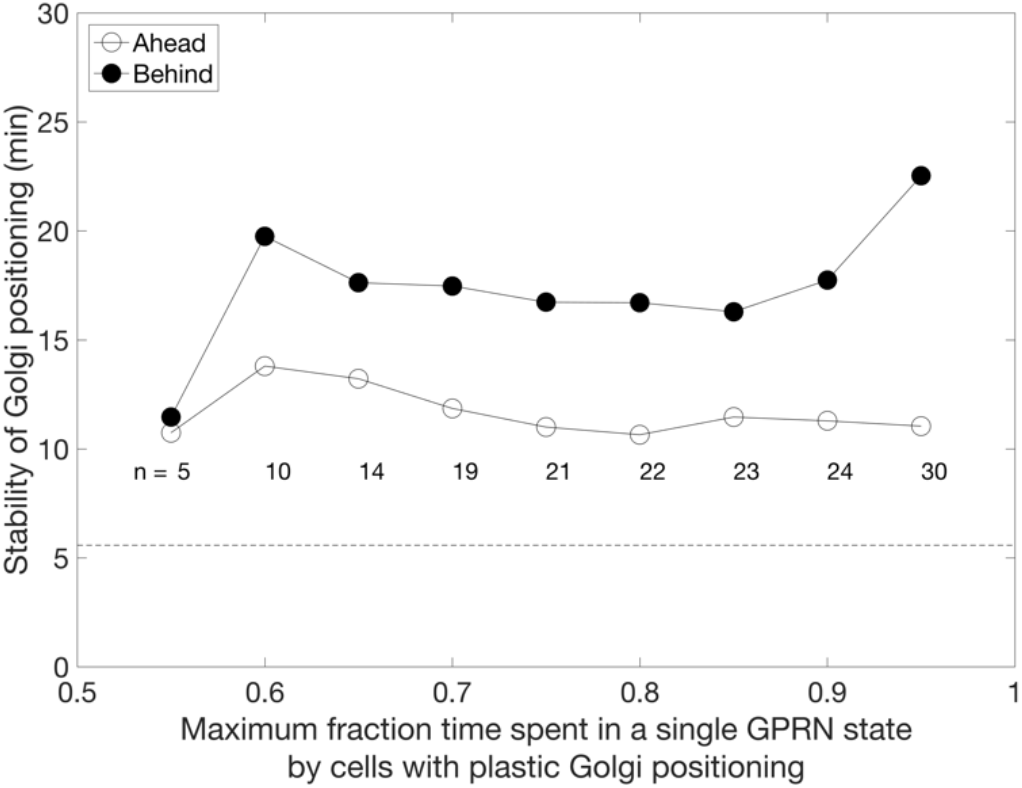
Golgi position is stabilized even among TGFβ-responsive cells with plastic Golgi positioning. The maximum fraction of time that a cell keeps its Golgi in a single position and still be categorized as having plastic Golgi positioning was varied (x-axis). The number (n) of TGFβ-responsive cells that meet the plasticity criterion is indicated. For each level of plasticity, the stability of Golgi ahead (open circles) and behind (filled) states was determined among plastic TGFβ-responsive cells. For comparison, the dashed line marks Golgi positional stability among TGFβ-treated, non-responsive cells.

At each level of the plasticity criterion, we identified plastic cells and quantified the stability of GPRN states A and B (Figure 7). Excluding cells that spend more than 90% of time in state A or B, the remaining plastic cells (n=24 cells) exhibit an average stability of 11.3 and 17.7 min for states A and B, respectively — stabilities that are 2.4- and 2.9-fold greater than those among non-responsive cells. Looking at cells that are more plastic and split their time at least 80:20 in one state versus another (n=22 cells), the average stability of states A and B were 10.6 and 16.7 min, values that are 2.1- and 2.7-fold greater than those among non-responsive cells. Even at a plasticity of 60:40 (n=10 cells), the stabilities of the A and B states (13.8 and 19.8 min) remain considerably higher (3- and 3.3-fold) than among non-responsive cells, albeit with the expected reduction in the sample size of cells that meet the plasticity criterion. These results further confirm that even among the most plastic TGFβ-responsive cells, Golgi states are significantly stabilized compared to non-responsive counterparts.

In summary, our analysis shows that TGFβ-responsive cells with enhanced motility on fiber-like micropatterns exhibit considerable stabilization of Golgi positioning. The stabilization is strong even among cells that exhibit plasticity in Golgi positioning; meanwhile, a non-negligible 35% of the population achieve complete stabilization and commit in an all-or-none manner to always maintaining the Golgi ahead or behind the nucleus. Our data shows that this stabilization effect occurs for both the Golgi ahead and behind states, albeit the effect is stronger for state B.

### Stability of Golgi positioning, irrespective of ahead or behind the nucleus, corresponds to singlecell migration efficiency

The data show that the TGFβ-responsive cells significantly stabilize Golgi positioning, both ahead and behind the nucleus. At the population level, the stabilization of GPRN corresponds to enhanced motility relative to the untreated and non-responsive cell populations. We next asked whether this relationship between GPRN stability and cell migration behavior applies even at the level of individual TGFβ-responsive cells.

To address this question, we used Pearson’s correlation to analyze to what extent variations in the stability of Golgi state helps to explain variations in migration speed and persistence of individual TGFβ-responsive cells, and whether this relationship is stronger for state A versus B. The stabilities of both states A and B exhibit a statistically-significant correlation with speed (Figure 8). Variations in the stability of states A and B capture approximately 22% and 30% of the variability in migration speed, respectively. Furthermore, the Spearman correlation indicates a statistically-significant monotonic relationship between cell migration speed and the stability of both Golgi states (data not shown, p < 0.01). Similarly, the stability of both states A and B exhibit a statistically-significant correlation with the persistence of migration and capture approximately 33% and 15%, respectively, of the variability in persistence. This analysis demonstrates that stable maintenance of Golgi position – either ahead or behind – the nucleus is associated with enhanced cell motility at the single-cell level.

**Figure 8.**
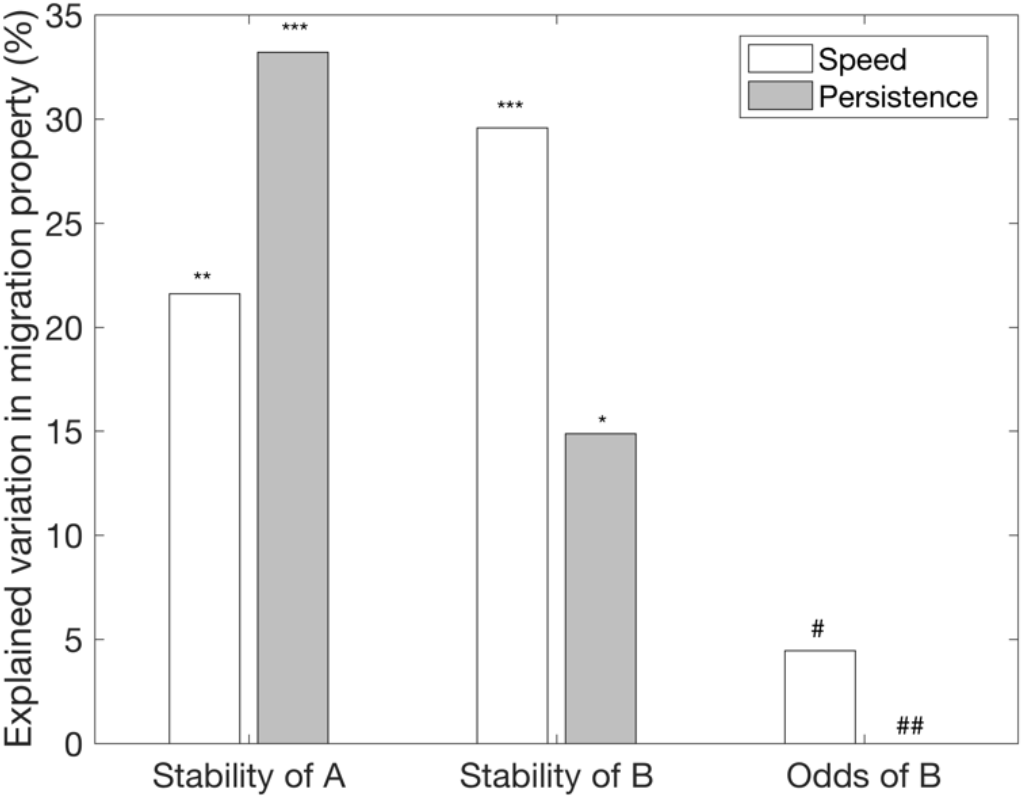
The stability of Golgi positioning — but not a bias for one position over another — exhibits a statistically-significant relationship to single-cell migration properties. The square of the Pearson’s correlation coefficient (PCC) was used to determine the degree to which singlecell variations in the stability and odds of positioning the Golgi ahead or behind the nucleus explains variations in migration speed (open) and persistence (filled) among TGFβ-responsive cells. The PCCs for relating migration properties to stability of Golgi position are statistically significant (***, p < 0.01; **, p < 0.01; * p < 0.05), whereas PCCs for relating migration behavior to the odds of finding the Golgi behind the nucleus are statistically insignificant (#, p > 0.2; ##, p > 0.95).

In contrast to the observed relationship between cell motility and the stability of GPRN states, the data show poor correlation between cell migration properties and the relative abundance or the odds of state B over state A (Figure 8). Thus, the relative abundance of time that a cell spends in one state over the other is not strongly linked to migration performance.

## DISCUSSION

We report here a novel relationship between Golgi positioning and cell motility. We show that the stabilization of Golgi positioning with respect to the nucleus — independent of whether the Golgi is ahead or behind the nucleus — corresponds to enhanced migration speed and persistence. TGFβ-mediated EMT stabilizes Golgi positioning by 3-4 fold while doubling and quadrupling speed and persistence of cell migration along fiber-like micropatterns. Furthermore, single-cell variations in Golgi stability capture up to 30% of variations in speed and persistence, whereas the bias in Golgi positioning is not statistically predictive of cell migration behavior. Since EMT and the development of a fibrillar microenvironment are significant cell-intrinsic and extrinsic drivers of breast cancer progression (3, 31), our findings have implications for understanding and therapeutically-targeting cell migration and invasion in cancer.

### EMT enhances motility in an already motogoenic fibrillar microenvironment, with implications for invasiveness during cancer progression

Here, we show that TGFβ-induced EMT enhances cell migration along fiber-like tracks compared to untreated epithelial cells. That EMT enhances motility is well-established in 2D contexts. That EMT also enhances migration in spatially-confined fiber-like environments is significant in several ways. First, it shows that cell-intrinsic progression through EMT and fiber maturation in the tumor microenvironment are non-redundant pathways that have the potential to cooperate to enhance motility during cancer progression. Second is the magnitude of the EMT effect: migration speed doubles and persistence quadruples in response to TGFβ-mediated EMT within spatially-confined tracks. These effects are on top of the well-documented positive effects that fibrillar topology alone has on cell migration of non-transformed epithelial cells when compared to non-fibrillar 2D and 3D microenvironments (13, 16, 17).

Third, the combined effect of EMT and fibrillar maturation on motility is seen not only at the level of single-cell motility, but also at the level of migration response to pairwise cell-cell interactions. We recently showed that when non-transformed epithelial cells encounter another cell along a fiber-like track, they reverse direction (11, 15). Progression through TGFβ-mediated EMT shifts this behavior, enabling cells to slide past each other and continue to move in their original direction. In a tumor environment crowded with cells, gaining the ability to slide allows cells to maintain their migration direction and achieve more effective dispersion, whereas cells that reverse direction at every cell-cell encounter will disperse more slowly.

Thus, the emerging picture is that EMT and fibrillar maturation in the microenvironment cooperate at the levels of individual cells and on cell-to-cell interactions to affect cell migration. A better understanding of this hierarchical cooperation will provide a foundation for predictive multiscale, multicellular models of dispersion and invasion in the tumor microenvironment (32).

### A role for stable Golgi positioning in EMT-mediated enhancement of cell motility in fibrillar contexts

How does EMT enhance migration along spatially-confined fiber-like tracks? The data indicate a role for the stabilization of Golgi positioning with respect to the nucleus (GPRN). Cells that respond to TGFβ with enhanced motility also stably maintain GPRN, with the Golgi either ahead (A) or behind (B) the nucleus, for 3-4 fold longer duration than cells that are untreated or are non-responsive to TGFβ. In fact, the stabilization is complete in approximately 35% of TGFβ-responsive cells that spend all their time in either the A or B state. Furthermore, among the remaining 65% TGFβ-responsive cells in which Golgi positioning is plastic, shifting between A and B states, the lifetime of each state is much greater than that observed among untreated or non-responsive cells.

Physical confinement along fiber-like tracks is, by itself, insufficient to stabilize Golgi positioning. Untreated non-transformed epithelial cells have relatively unstable GPRN compared to TGFβ-treated cells. Thus, although spatial confinement is sufficient to confer a uniaxial cell morphology and enhance motility compared 2D environments (13), EMT provides additional cell-intrinsic changes that stabilize Golgi position and promotes a 4-fold increase in persistence relative to cells that have not undergone EMT.

In fact, single-cell variations in the stability of Golgi positioning is sufficient to capture up to 30% of the variability in migration speed and persistence among TGFβ-responsive cells. This level of correspondence is remarkable given the many physical and molecular mechanisms involved in a complex behavior, such as cell migration. In contrast to the positional stability of the Golgi, the bias in positioning the Golgi behind the nucleus is statistically insignificant in explaining variation in cell migration behavior.

In future work, it is reasonable to hypothesize that combining additional molecular and physical properties — e.g., nuclear shape and deformation, properties of focal adhesions, etc. — with the stability of Golgi positioning could yield a multifactorial, predictive model of cell migration dynamics, akin to systems-level models of apoptosis composed of MAP kinase predictors (33). The inclusion of physical properties, such as Golgi stabilization, as predictors is consistent with the physiochemical nature of the cell migration process.

### The role of Golgi in migration is independent of its position relative to the nucleus

Our data indicate that the role of the Golgi in EMT-mediated enhanced fibrillar migration is independent of whether it is ahead or behind the nucleus. Whether a cell spends more time in state B than in state A has less than 5% association with enhanced migration. In contrast, the stabilities of both the A and B states are more strongly associated with enhanced speed and persistence of TGFβ-responsive cells. Furthermore, treatment with TGFβ enhances cell migration and stabilizes both A and B states, indicating that overall positional stability is more important than stabilizing one state preferentially. Thus, although TGFβ-responsive cells exhibit a population-averaged 2:1 bias in the prevalence of GPRN state B versus A, single-cell analysis shows that stability of Golgi positioning, more so than the actual position of the Golgi, is quantitatively related to cell migration behavior.

The most developed paradigm of Golgi positional bias in migration is based on 2D wound healing models in which the Golgi, together with the MTOC/centrosome, is positioned ahead of the nucleus from where it traffics proteins to the leading edge, resupplying molecular components, such as integrins, necessary for adhesion and traction (20–22). The Golgi has also been shown to nucleate its own set of microtubules that are preferentially oriented toward the leading edge, potentially providing avenues for polarized trafficking (34, 35).

However, there are many exceptions to anterior Golgi positioning. Even during wound healing in 2D, centrosome positioning — and by inference, Golgi position — is cell type-specific, with anterior positioning in CHO cells and posterior positioning in PtK cells (36). Meanwhile, chicken embryo fibroblasts exhibit anterior positioning of the centrosome on 2D glass, but exhibit no bias when migrating in collagen gels and along micron-scale grooves (37). Indeed, among individual Rat2 fibroblasts, no correlation is observed between Golgi orientation and migration speed on 2D surfaces (24).

Furthermore, rearward bias in Golgi positioning is observed in cells migrating on narrow micropatterns. African green monkey kidney cells (Bsc1) position the Golgi posteriorly 70% of the time during migration along 6-14 μm micropatterns (12). Another group observed rearward bias in 3T3 fibroblasts, albeit less pronounced (13). Meanwhile, we report here no bias in untreated mammary epithelial cells and a 60% rearward bias in TGFβ-treated cells migrating along 10 μm fiber-like micropatterns.

Considering these observations altogether, it is evident that cells are capable of migrating with anterior, posterior and even no bias in Golgi position in a context- and cell type-dependent manner. Furthermore, in some situations, such as the aforementioned studies in fiber-like contexts, the positional bias is quantitative and not qualitatively exclusive, suggesting that anterior vs posterior positioning of the Golgi is not an overriding factor in motility in these systems. Meanwhile, our findings point to a significant role for the stability of Golgi position in fibrillar migration, and in future studies, it will be important to test the relationship between Golgi positional stability and motility across a wide range of cell types and contexts.

### A working hypothesis for a structural/physical role for the Golgi in cell motility

How is stable positioning of the Golgi — independent of whether it is ahead or behind the nucleus — advantageous to migration along fiber-like tracks? We propose a structural and physical role for the Golgi in cell motility. The Golgi is part of a multi-component intracellular scaffold. It physically interacts and associates with microtubules, the microtubule organizing center (MTOC) and the nucleus (19). The Golgi is also associated with the actin cytoskeleton (38, 39), and particularly relevant in the context of EMT, it interacts with the vimentin intermediate filament network (40). We hypothesize that the Golgi, as a component of this intracellular scaffold, bears load from cell-generated forces and affects the transmission of these forces across the cell. If the Golgi position undergoes frequent changes, its effect on the spatial distribution of forces at one instant in time could be counteracted by contributing to a different, conflicting force distribution later in time. To the extent that the Golgi position can be maintained, its role in distributing forces is consistent over time, and migration is made more efficient.

Additionally, a canonical arrangement of the Golgi next to the MTOC next to the nucleus is well known (19). Every change in Golgi position could disrupt this three-member structure, and the rate at which the Golgi-MTOC-nucleus arrangement is reacquired may limit motility. Increased Golgi positional stability would minimize this disruption.

The extent to which the Golgi regulates the assembly and maintenance of an intracellular “core” scaffold mediating migration is unclear. Most likely, the role of the Golgi must be understood in the context of an integrative, systems-level crosstalk among organelles and cytoskeletal components. Whether stable Golgi positioning regulates, or is an indicator of, a stable migration-promoting core scaffold will help determine whether it could be targeted as part of a therapeutic strategy or be utilized in an imaging-based drug screening platform, respectively. In either case, the reported findings and implications underscore a need to more deeply understand the regulatory mechanisms and downstream consequences of stable Golgi positioning during cell migration.

## CONCLUSIONS

EMT and maturation of collagen fibrils in the microenvironment are associated with cancer progression. To the extent that these factors can act as non-redundant promoters of motility, their simultaneous occurrence during cancer progression has the potential to accelerate motility and invasion. In this context, we find that EMT significantly enhances migration in a spatially confined, 1d microenvironment that is already known to be more conducive to motility than an unconstrained 2d context. Moreover, single-cell analysis shows that EMT-mediated stabilization of Golgi position, and not a preference for anterior or posterior positioning, is associated with enhancements in migration speed and persistence. These results suggest a model in which the Golgi is part of a core physical scaffold whose stability provides a reliable platform for coordinating and transducing forces across the cell, thereby enabling more efficient migration.

## SUPPORTING MATERIAL

Supporting video is included with the submission.

## AUTHOR CONTRIBUTIONS

RJN and ARA conceived and designed the experiments. RJN performed the experiments. RJN and MLL conducted the image analysis. RJN and ARA analyzed results and wrote the manuscript. All authors edited the manuscript.

## ACKNOWLEDGEMENTS

This work was funded by NIH grant R01CA138899 to ARA, NIH grant R01CA098830 and the Era of Hope Scholar Award from the DOD Breast Cancer Research Program to SKM. We thank Nick Ngai for making and sharing the MCF-10A cell line expressing nuclear and Golgi markers.

